# Nodule branching, size, and symbiosis outcomes shaped by natural genetic variation in rhizobia and alfalfa

**DOI:** 10.1101/2025.10.27.684485

**Authors:** Elizabeth L. Paillan, Alejandra Gil-Polo, Sohini Guha, Andy Swartley, Liana T. Burghardt

**Affiliations:** The Pennsylvania State University, Department of Plant Science, University Park, Pennsylvania; North Carolina State University, Department of Plant and Microbial Biology, Raleigh, North Carolina

**Keywords:** Bacterial fitness, Legume-rhizobia mutualism, Medicago sativa, Nitrogen-fixation, Nodule morphology, Partner quality, Sinorhizobium meliloti

## Abstract

In many host-microbe symbioses, hosts develop specialized organs that house microbial symbionts. In the legume-rhizobium symbiosis, these organs are called root nodules. While many studies have focused on host traits like plant growth, nodule number, and their structural changes, nodule morphology may also influence rhizobial benefits (e.g., bacterial population size). Here, we investigated how variation in alfalfa nodule morphologies influences the fitness outcome of both partners. Specifically, we measured how host and strain genetic variation influenced 1) nodule branch number, area, and rhizobial fitness at the individual nodule level and 2) the proportion of branched nodules, nodule number, and plant benefits at the whole plant level. From the rhizobial perspective, host identity had a stronger effect on rhizobial fitness, and larger or more branched nodules tend to release more rhizobia. From the host perspective, the proportion of branched nodules was strongly negatively correlated with the total nodule number and positively correlated with host benefit across multiple alfalfa genotypes. However, the strength of the relationship differed among hosts. Together, these results provide insight into the host and rhizobial genetic contributions to nodule morphology and symbiotic outcomes, suggesting a novel direction for further research in legume-rhizobium mutualism.

**Highlight:** Plant and bacterial symbiotic effectiveness can be gauged by the number of nodule branches formed in indeterminant legumes-rhizobium systems along with traditional metrics such as nodule count.

## Introduction

In host-microbe symbiosis, microbes often reside in host tissues within chimeric organs (Moran, 2007; Oldroyd et al., 2011; Visick et al., 2021). These organs, thus, create the environment that shapes microbial fitness outcomes. In the rhizobia-legume symbiosis, these chimeric organs are known as nodules, which form on the roots of legumes (Oldroyd, 2013). Nodules can exhibit diverse shapes; yet we know little about the factors contributing to the various morphologies or consequences these morphologies have on benefits that hosts and microbes receive during symbiosis (Dupont et al., 2012; Sachs et al., 2018). More specifically, at present, little is known about how nodule branching in indeterminate nodules influences rhizobial fitness and host fitness, and whether nodule branching is driven by host or rhizobial identity (Lindström and Mousavi, 2019). Here, we address these questions by examining multiple hosts and rhizobial genotypes to disentangle their contributions towards nodule branching and how it affects rhizobial and host fitness.

Legume root nodules follow two developmental patterns broadly categorized as determinate or indeterminate based on the meristematic behavior of the host plant. Determinate nodules, found in soybean (*Glycine max*) and common bean (*Phaseolus vulgaris)*, lack a persistent meristem, taking on a spherical shape. In contrast, indeterminate nodules, the focus of this investigation, found in species such as barrel clover (*Medicago truncatula)* and its close relative alfalfa *(Medicago sativa)*, can take an elongated shape due to a persistent nodule meristem that sustains prolonged growth (Ferguson et al., 2010; Sprent et al., 2017). Indeterminate nodules branch when the nodule meristem splits after the median part of the nodule meristem stops dividing, leading to multiple branches, each one with its own meristem (Łotocka et al., 2012). Therefore, depending on the branching level, indeterminate nodules encompass a broad range from uni-branched and bifurcated to palmate and coralloid structures (Hirsch, 1992; Łotocka et al., 2012; Gola, 2014; Cai et al., 2018). To our knowledge, the effect of nodule branching on the fitness of rhizobia and plants has yet to be investigated.

In successful rhizobia-legume symbiotic associations, both partners enhance each other’s fitness (Denison and Kiers, 2004). Rhizobia convert inaccessible atmospheric nitrogen into an accessible form used for plant growth. In return, the plant provides rhizobia with sugars derived from photosynthesis, which the rhizobia will use to grow and divide inside the nodules (Oldroyd, 2013). Within indeterminate nodules, rhizobia segregate into two zones: (1) terminally differentiated, nitrogen-fixing bacteroids encapsulated by a plant-derived membrane, and (2) an undifferentiated rhizobial population (Thies et al., 1991; Sachs et al., 2018). The bacteroids concentrate in the nitrogen-fixing zone of the nodule, while undifferentiated bacteria live in the “infection zone” and “saprophytic zone” of the nodule (Vasse et al., 1990; Batut and Boistard, 1994; Timmers et al., 2000; Łotocka et al., 2012). These undifferentiated cells retain the ability to reproduce and are released back into the soil when nodules senesce; thus, the size of the undifferentiated rhizobia population in nodules is of particular interest when considering rhizobial benefits from symbiosis (Denison and Kiers, 2004). Branching morphogenesis could influence symbiotic outcomes by maximizing nodule area and overcoming spatial constraints imposed by organ size. Research suggests that the size of a root nodule can influence the rhizobial population it releases and, consequently, the rhizobial fitness (Lu and Werb, 2008; Burghardt, 2020) and because some legumes can allocate more resources to nodules harboring effective nitrogen-fixing strains, the nodule size can be positively correlated with plant fitness (Westhoek et al., 2021). Thus, nodule branching could affect rhizobia and plant fitness, respectively, and this could suggest new mechanisms by which rhizobia and hosts alter each other’s fitness (Porter et al., 2024).

Very little is known about how host or rhizobial genetics contribute to nodule branching and size. Some host genes have been characterized as being involved in nodule branching. For example, the overexpression or loss-of-function of NOD FACTOR HYDROLASE1 (MtNFH1) in *M. truncatula* and the mutation of the *cochleata* gene (*coch*) in *Pisum sativum* lead to increased nodule branching (Ferguson and Reid, 2005; Cai et al., 2018). On the rhizobial side, multiple genes have been identified as playing a role in controlling nodule morphology. In *Sinorhizobium meliloti*, overproduction of Nod factors within the nodule leads to increased nodule branching (Cai et al., 2018). Rhizobia are also speculated to affect nodule morphology through altered expression of phytohormone production. For example, *Rhizobium leguminusarum viciae* that overproduce indole-3-acetic acid induce larger nodules with a more active meristem, resulting in meristem splitting in *Vicia hirsute* (Camerini et al., 2008). Other host and rhizobial genes may well influence nodule branching, but it is not a commonly reported phenotype. Clearly, there is much more to be discovered about how the complex crosstalk between the host and rhizobia affects nodule branching.

Here, we explore the effects of host and isolate identity on nodule branching and rhizobial populations using natural genetic diversity of both partners. We used a collection of 117 *Sinorhizobium* isolates recently isolated from an alfalfa variety trial and performed a single-isolate inoculation experiment on three alfalfa (*M. sativa)* varieties in a controlled greenhouse to investigate the effect of nodule branching on rhizobial fitness. First, we sampled individual nodules from plants inoculated with a subset of eight strains. The nodules spanned a wide range of sizes and numbers of branches. We investigate whether nodule size and the number of rhizobia released from nodules are related to nodule branching, and if so, whether these relationships depend on the host variety or the rhizobium isolate identity. Second, to investigate the effect of nodule branching on plant benefits, we measured per-plant traits for all plants inoculated with each of the 117 strains. We investigated whether host variety or rhizobial isolate influences nodule branching propensity, and whether strains that cause more branched nodules also tend to have increased nitrogen-fixing efficiency.

## Materials & Methods

### Source of *Sinorhizobium meliloti* isolates

We developed a collection of 117 *Sinorhizobium meliloti* isolates from Alfalfa (varieties: Oneida, Vernal, and SW402) planted in 2016 at Penn-State’s Russell E Larson Agricultural Station in Rocksprings, PA (40.7215,−77.9275). Full details of this collection will be published separately, along with genome sequencing information (Guha et al. in prep). Briefly, in the fall of 2020, we sampled four-year-old alfalfa plants in 0.914 m × 6.019 m strips using a randomized block design. We dug up 80 nodules from each of the three varieties growing within 5 meters of each other. After surface-sterilizing the nodules and homogenizing them in a 0.85% NaCl solution, the Isolation Bio Prospector system was used to culture individual rhizobial cells. To further purify the isolates, we streaked individual cultures from each well on a Yeast Mannitol Agar plate, picked a single colony, re-streaked it, and incubated the plates at 28 °C (Vincent, 1970).

### Plant preparation for the single-strain inoculation experiment

To examine host-rhizobia interactions, we grew each isolate with four replicates of the three alfalfa varieties from the field under nitrogen-limited conditions. We sterilized 1,200 alfalfa seeds of each variety in 10% commercial bleach for 90 seconds, rinsed them with deionized (DI) water, and transferred them onto filter paper in glass Petri dishes. We wrapped the petri dishes in aluminum foil and placed them at 4 °C for three days to stratify. In the greenhouse, we washed and bleached Heavyweight Deepot™ Cells (D40H – 2.5” diameter, 10” depth; item number: D40H), (N=1440; 4 replicates x 3 alfalfa varieties x 120 isolates & controls), filled them with a mix of 50:50 sand and calcined clay. The pots were held in Deepot™ Support Tray (D20T for 2.5” & 2.7”; item number: D20T), arranged with an empty cell between each pot, and kept in four blocks. We planted two seeds in each pot, covered them with ½ to ¼ inch of sterile vermiculite, kept pots moist with twice-daily misting, and maintained humidity during germination by covering them with a clear plastic tarp. After one week, pots were moved to their randomized locations, and the plastic tarp was removed.

### Plant inoculation and care

*S. meliloti* isolates were streaked onto Yeast Mannitol agar plates from glycerol stocks stored at –80°C, placed in an incubator, and grown for four days at 28°C. Then, the bacteria were transferred to Tryptone Mannitol liquid media and grown overnight in a shaking incubator at 150 RPM at 28°C. We measured and adjusted each culture’s optical density (OD600) to reach 0.1, then diluted it to 50ml with DI water. Later, we inoculated each pot eight days post-sowing with ~ 10^6 CFU. We based the OD600-to-colony-forming unit correlations on correlations made with eight strains from our collection. Greenhouse conditions were maintained at 18°C to 23°C during the daytime and a minimum temperature of 12°C throughout the experiment. Three weeks post-inoculation, we began fertilizing alfalfa weekly with 10 mL of 2x Fahräeus medium from the *M. truncatula handbook Appendix 2* (Barker et al., 2006). We doubled the fertilizer concentration every two weeks to track plant growth. We thinned to one plant per pot four weeks after sowing using sterile scissors and tweezers. Each pot was watered individually as needed to prevent cross-contamination of inoculated isolates.

### Measurement of plant-level benefit traits

Eight weeks post-inoculation, we harvested plants and assessed above- and below-ground plant traits. First, we collected the aboveground biomass and dried it at 75°C for at least 72 hours. Second, we gently removed the roots from the growing media and briefly suspended them in water to remove any sand and calcined clay. We wrapped cleaned roots in a wet paper towel and stored them in a Ziplock bag at 4°C for up to five days to allow us to manually count the number of branched and unbranched nodules. After recording the nodule information, we dried the roots, just as we did the shoots. We assessed biomass on a VWR® P2 balance (VWR-203P2/C).

### Measurement of rhizobial fitness in individual nodules on a subset of host x strain combinations

During the above harvest, we collected six nodules from seven rhizobial isolates by three host variety combinations. The nodules were chosen, not randomly, but to span the nodule size variation on each plant. We photographed nodules on a green background to enable nodule area measurement and branch counting using the ImageJ “analyze particle tool” (Schneider et al., 2012). We then sterilized individual nodules with 20% commercial bleach and 5μl Polysorbate 20 for two to four minutes inside 1.7 mL Eppendorf tubes, lightly tapping the tube to ensure contact with all the nodule surface area. Each nodule was rinsed thrice with sterile water and stored in 100μL of 0.85% NaCl at 4 °C overnight. Using a sterile pestle, we crushed the nodules, added 900 μL 0.85% NaCl to reach 1mL, and performed a dilution series to determine the number of undifferentiated bacteria in each nodule. For smaller nodules, we plated and counted five 20 μL spots for both the 10^3^ and 10^4^ dilutions, while for larger nodules, we spot-plated the 10^5^ and 10^6^ dilutions. Spots were counted ~48 hours after treatment and then double-checked at 72 hours. We determined the dilution level at which the number of colonies mostly ranged between 10 and 100 and calculated the average number of colonies across all five spots. We multiplied the average by the dilution factor and divided by the volume plated to estimate the population size of rhizobia released from each nodule.

### Statistical Analysis

All statistical analyses were performed using the R statistical environment (version 4.4.3; R Core Team, 2016). To test the relationships between single nodule area, rhizobial population size, and the number of branches, we fit linear models and tested for significant terms using an analysis of variance (ANOVA). To meet a normal distribution assumption, we used a log10 transformation of our nodule surface area measurements and bacterial colony-forming units (CFU) measurements. We used the following models: To test if nodule branching increased nodule surface area across host varieties and rhizobial isolates we used the model *log*_*10*_ *Nodule Surface area ~ number of nodule branches * Alfalfa host variety * rhizobial isolate + replicate;* To assess how the number of nodule branches influenced the undifferentiated *Sinorhizobial* population, we used the model *log*_*10*_ *Rhizobial bacterial colonies ~ Number of branches* Alfalfa host variety + Rhizobial isolate + Replicate*. To test if the number of branches altered the relationship between nodule size and rhizobial bacteria released from nodules, we used the following model: *log*_*10*_*(Rhizobial bacteria colonies)~ Nodule surface area* Number of branches Alfalfa host variety + Rhizobial isolate + Replicate*.

To assess the relationship between our measured predictors of plant benefit, we used the proportion of branched nodules (number of branched nodules / (number of unbranched nodules + number of branched nodules)) and host biomass (aboveground + belowground biomass). We fit linear models and tested for significance using an ANOVA. We used the following models: To assess if plants with a higher proportion of branched nodules tended to form more nodules while accounting for rhizobial strain variation *(Mean nodule count ~ mean proportion of branched nodules + rhizobial isolate);* To assess if strains that produce a higher proportion of branched nodules also support more host growth and if the effect depend on host variety we ran the following model mean (*host biomass~ mean proportion of branched nodules * Alfalfa host variety + rhizobial isolate)*. To test the contributions of host and strain genetics to nodule branching propensity, we ran the model *(proportion of branched nodules ~ Alfalfa host variety *rhizobial isolate + replicate)*.

## Results

### Bigger and more branched nodules have larger populations of reproductively viable rhizobia, and host genetics plays a larger role than rhizobial genetics

First, we tested if nodule branching and size covary and determined if the relationship is influenced by host and isolate genetics (Table S1, *R*^*2*^_*adj*_ *=0*.*6678*). We found, perhaps unsurprisingly, that nodules with more branches tended to have a larger surface area (*p* = *4*.*14×10*^*−48*^). Host genotype did not significantly alter the relationship between these traits (*p =* 0.607), whereas isolate identity did (*p =* 0.001, Fig. 2a). This implies that some strains tend to form smaller or larger nodules, independent of nodule branching and host identity. In summary, nodule branching strongly correlates with nodule surface area, and this relationship is predominantly influenced by isolate identity, rather than the host. Next, we examined the relationship between rhizobial population size (inferred by the number of undifferentiated *Sinorhizobium* released *from* each nodule) and the number of nodule branches (Figure 2b, Table S2, full model *R*^*2*^_*adj*_ *=0*.*2116*). We found that the population of *Sinorhizobium* increases with the number of nodule branches (*p=4*.*66×10*^*−11*^), and that this relationship is somewhat dependent on host variety (*p*=0.002) but not on strain identity (*p*=0.34). While some unbranched nodules also house large culturable rhizobial populations, more branched nodules consistently release large populations of undifferentiated *Sinorhizobium*. This supports our hypothesis that forming branched nodules increases rhizobial fitness by expanding the nodule region that undifferentiated rhizobia can inhabit (Fig. 2b).

**Figure 1:**
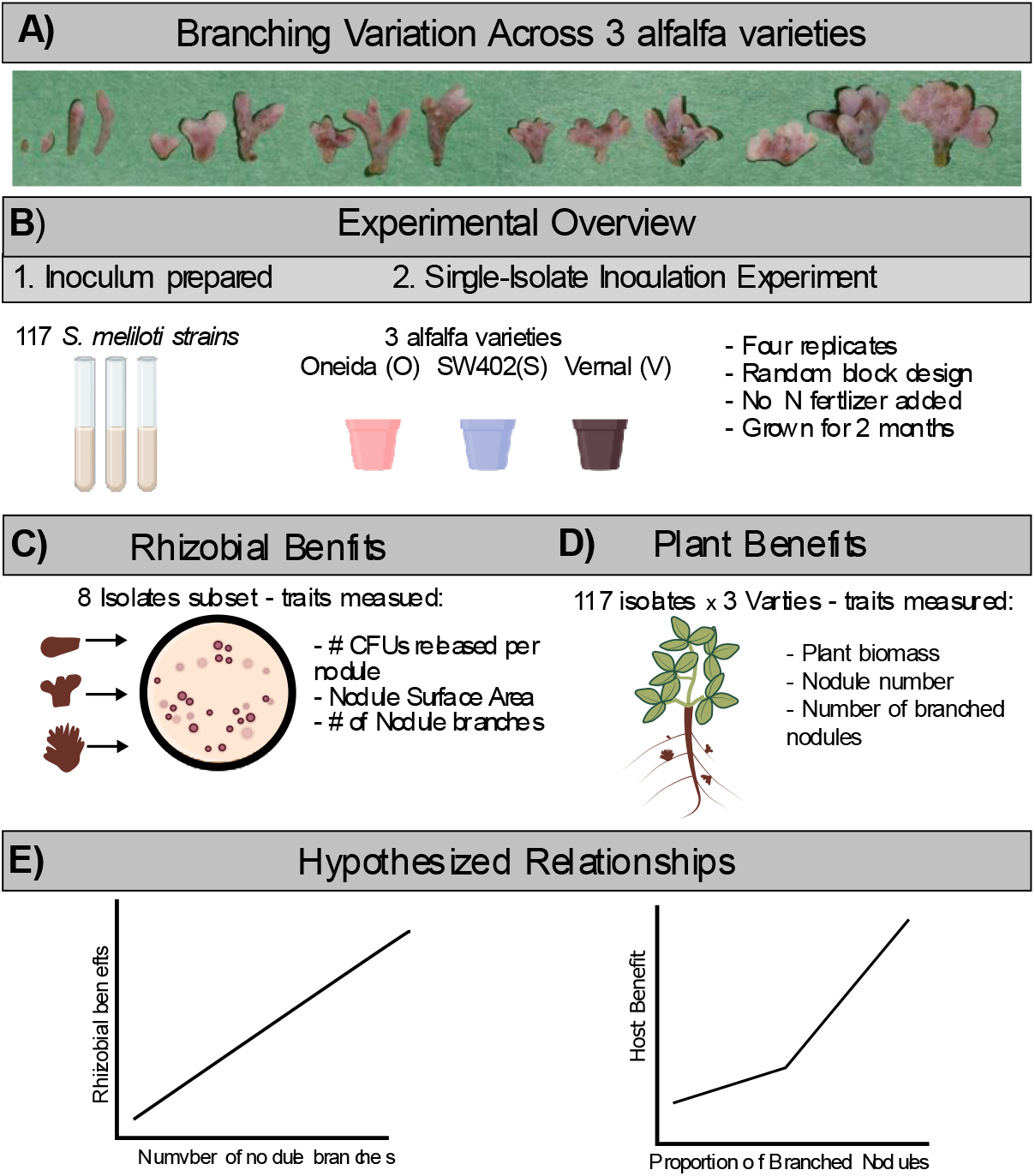
Overview of nodule branching, experimental design, and hypothesized relationships. **A)** Nodules ranging from 1 – 5+ branches respectively collected from inoculated alfalfa. **B)** Overview of our main experiment with 117 *Sinorhizobium* isolates inoculated in 3 genotypes of *Medicago sativa*, alfalfa. **C)** Measured rhizobial benefit traits from an 8 isolate subset from the main experiment. **D)** Measured plant benefit traits from all four replicates of 3 alfalfa varieties singly inoculated with 117 isolates. **E)** Our hypothesis of the tested relationships between rhizobial benefit traits and the number of nodule branches, and the relationship between host benefit and the proportion of branched nodules. Created in Biorender. Burghardt, L. (2025) https://BioRender.com/soj4e5a.

**Figure 2:**
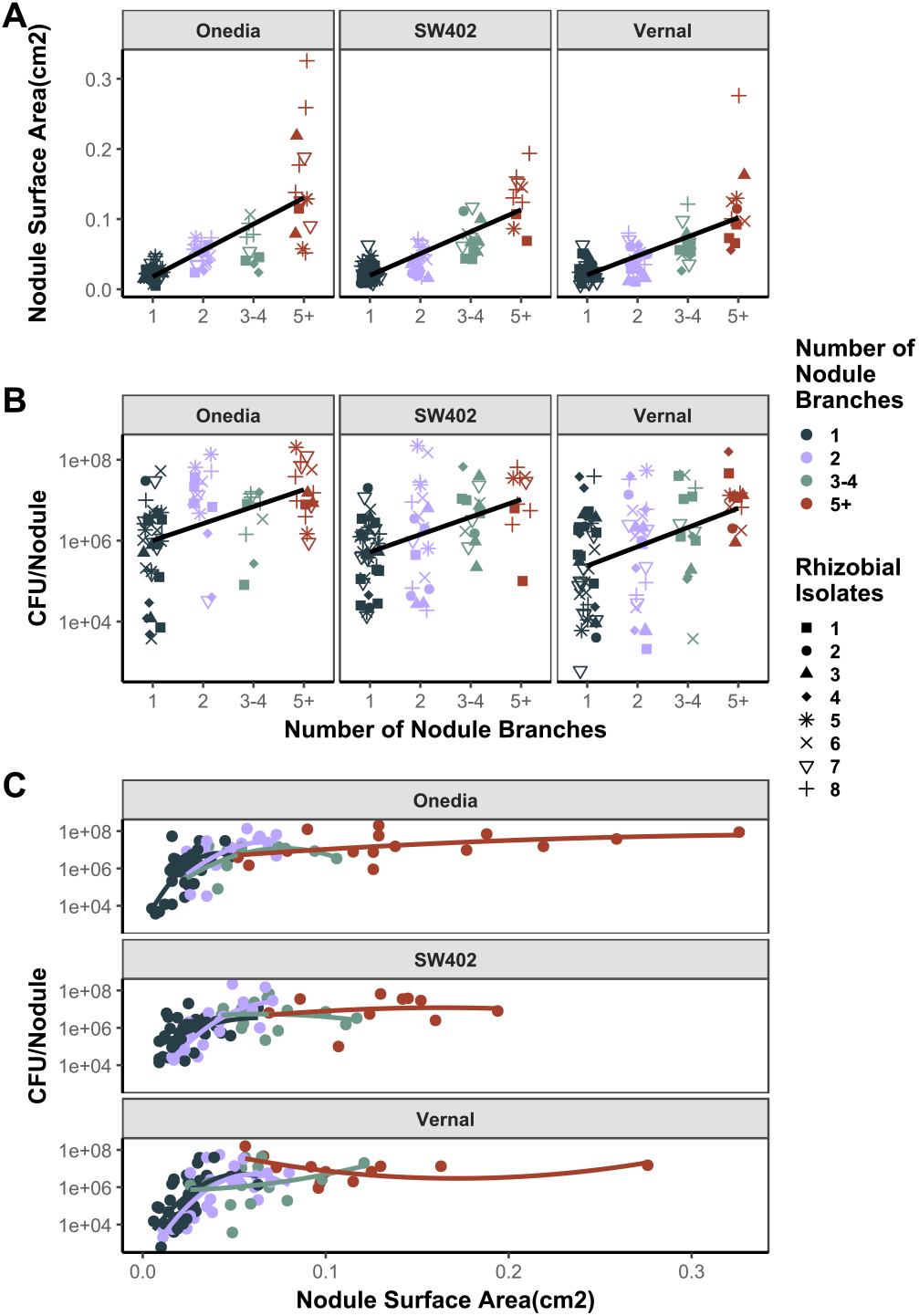
Relationships between individual nodule traits and rhizobia released across alfalfa host varieties *Oneida, SW402, and Vernal*. **(A)** Scatterplot of the nodule surface area across nodule branch number *(p= 4*.*14×10*^*−48*^, *R*^*2*^*=0*.*6678)*. **(B)** Scatter plot of the CFUs of *Sinorhizobium* released from crushing nodules across nodules with 1-5+ branches presented on a log_10_ scale (*p=4*.*66×10*^*−11*^, *R*^*2*^*= 0*.*2116)*. **(C)** Scatter plot of the CFUs of *Sinorhizobium* released from crushing nodules across nodules with 1-5+ branches present on a log_10_ scale (*p=1*.*50×10*^*−21*^, *R*^*2*^*=0*.*453*). The more branches a nodule has, the higher the total nodule surface area, and the number of *Sinorhizobium released* when crushed is increased. However, there is a limit to this effect. *Additional details: Anova and model details are in (Supplement Table 1)*.

**Figure 3:**
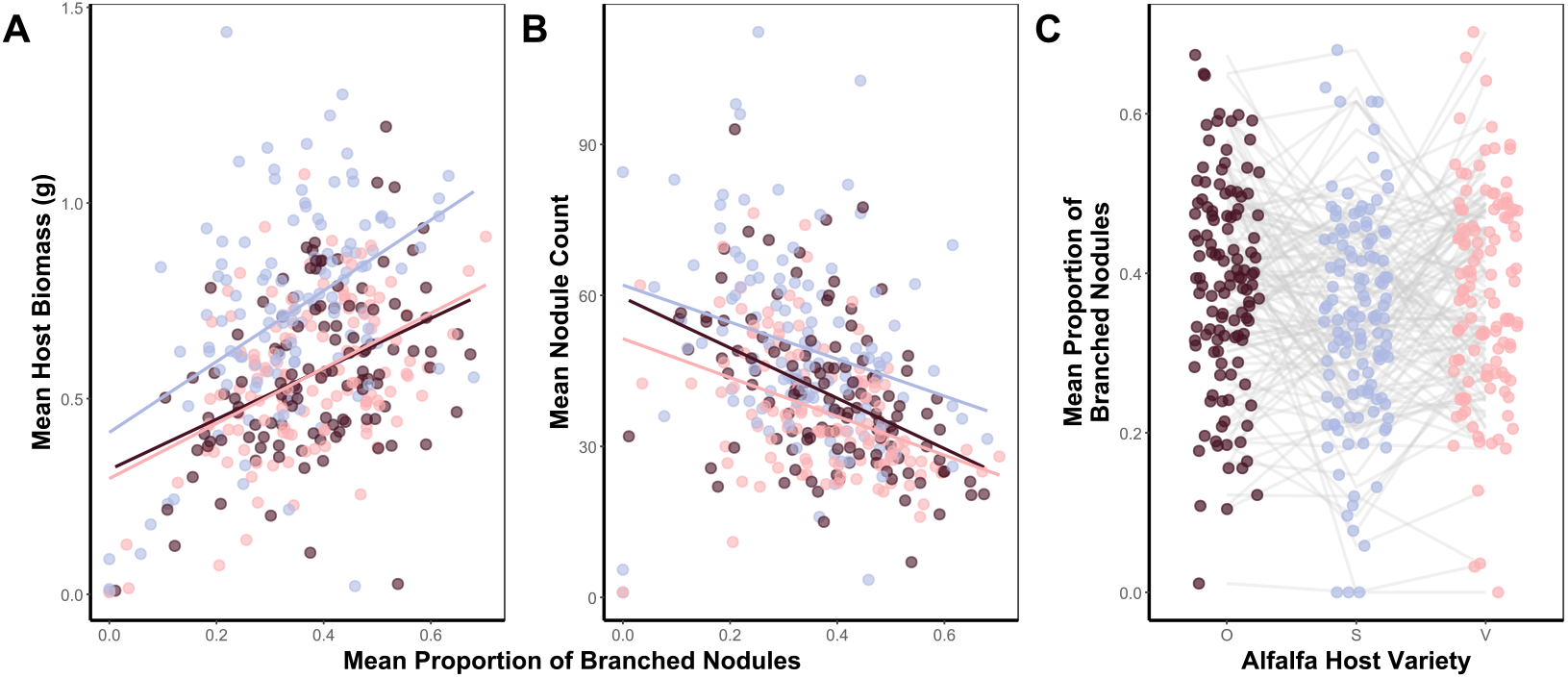
Relationship between nodule traits and plant benefits across alfalfa hosts, where each point represents the mean across four replicate pots. **(A)** Scatterplot of the mean host biomass across mean proportion of branched nodules. (*p=1*.*13×10*^*−22*^, *R*^*2*^*=0*.*530)*. **(B)** Mean nodule count across host plant-Sinorhizobium isolate combination replicates and its correlation to the mean proportion of branched nodules *(p=7*.*89×10*^*−17*^, *R*^*2*^*=0*.*458)*. Increasing the number of branched nodules can increase the host plant’s dried biomass while contributing to a lower total count of nodules overall. **(C)** *Sinorhizobium* isolate performance depends on the host and isolate genotype separately when forming branched nodules (p=*3*.*19 × 10*^*−20*^, *R*^*2*^*=0*.*454). Host alfalfa genotypes include Oneida, SW402, and Vernal. Additional details: Anova and model details are in (Supplement Table 2)*.

Our results also suggest that branching alters the relationship between nodule size and the culturable rhizobial population. Given that branch number and nodule area are correlated, we observed that rhizobial population sizes increased with nodule size (Fig. 2c). However, unlike nodule branch number, there is an apparent plateau, such that increasing nodule surface area does not lead to more culturable *Sinorhizobium* from nodules. This decoupling of nodule surface area and rhizobia population size could be due to the spatial restriction of infection threads near the meristematic region on the nodule branches. To explore this possibility, we ran a third model to explore if nodule branching influences the relationship between CFU released and nodule surface area (Table S1, *R*^*2*^_*adj*_ *=0*.*453*). We found that the relationship between nodule surface area and rhizobia released significantly changed depending on the number of branches a nodule had *(p = 1*.*50×10*^*−21*^). Nodule area and rhizobia release are tightly coupled when nodules have only one or two branches (Fig. 2c), the relationship erodes, and rhizobia population plateaus when nodule branching increases. Moreover, host varieties differed overall in rhizobia released per nodule independent of the relationship between size and branches (*p*=0.001, Fig. 2c). Overall, we find that nodules that have fewer than two branches and were relatively large, released a similar amount of rhizobia as nodules with three or more branches (Fig. 2c). Taken together, this suggests that size is not the sole driver of viable *Sinorhizobium* inhabiting a nodule, the architecture, branching, and host genetics play a role in applying a physical constraint to this relationship.

### Rhizobial isolates that produce fewer, more branched nodules with a host tend to provide more benefits

We extended our analysis to include the effect that nodule branching has on host traits, such as nodule number and total host biomass. We found evidence that a tradeoff exists between forming more nodules and producing a higher fraction of branched nodules (*p=1*.*13×10*^*−22*^), and that both host variety (*p*=*1*.*7×10*^*−20*^) and specific *Sinorhizobium* isolate (*p*=*1*.*27×10*^*−9*^) significantly altered this relationship (full model *R*^*2*^_*adj*_ *=0*.*530)* (Table S3). One possible interpretation is that branched nodules are involved in a regulatory strategy for the host, where increased investment in branching complexity reduces the total number of nodules produced. Building on previous results, we sought to determine if the nodule branching tendency of a *Sinorhizobium* isolate was associated with greater host benefits. We found that rhizobial isolates that produced a greater proportion of branched nodules with a host variety tended to be more beneficial (*p* = *7*.*89×10*^*−17*^) (*R*^*2*^_*adj*_ *=0*.*458*). This relationship depends on the host variety (*p = 3*.*11×10*^*−11*^*) and the Sinorhizobium isolate identity (p = 2*.*28×10*^*−9*^) separately (Table S3 for full model details). This suggests that *Sinorhizobium* isolates that produced more branched nodules tended to support larger host plants. Because the plants were grown in a nitrogen-free environment, this increased growth can be attributed to increased nitrogen fixation (Fig.3b).

## Discussion

In this investigation, we examined nodule branching from both the rhizobial fitness and the plant host fitness perspectives, focusing specifically on the effect of genetic variation in both the rhizobia and alfalfa. We found that larger, branched nodules release more rhizobial bacteria regardless of the host or isolate identity. From the plant host perspective, we found that the proportion of branched nodules can be used as a low-cost and low-effort proxy for the nitrogen-fixing benefits each strain provides to the host. More broadly, these results suggest a positive correlation between rhizobial fitness benefits (i.e., rhizobia released from the nodules) and plant benefits (i.e., fixed nitrogen). From an ecological perspective, our results support the hypothesis that host plant resource allocation may contribute to nodule branching. Below, we discuss the implications of these findings for 1) understanding rhizobial fitness, 2) using branching as a proxy of host benefit, 3) the contribution of host and rhizobial genetics to branching, and 4) maintaining mutualisms.

### Nodule branching alters the relationship between nodule size and rhizobial population size

To our knowledge, this is the first study on the effect of branching on rhizobial population size. Our results suggest that branching alters the structure of nodules, leading to larger rhizobial populations. This effect is likely connected to shifts in the relative areas of the five nodule zones that characterize indeterminate nodules and that differentially support rhizobial reproduction and nitrogen fixation (Roux et al., 2014). However, the increase in rhizobia released from nodules levels off beyond three branches, indicating a saturation point where further branching provides little additional benefit to rhizobia. The observed uncoupling could reflect multiple physiological ceilings, including oxygen diffusion limits (Tjepkema and Yocum, 1973), carbon supply constraints (Udvardi and Poole, 2013), developmentally regulated limits on nodule expansion (Ferguson et al., 2010; Mortier et al., 2012), and host sanctions that restrict undifferentiated rhizobial proliferation (Kiers et al., 2003; Oono & Denison, 2010). While our nodules were a maximum of 8 weeks old, our results suggest further avenues for research including measuring undifferentiated population sizes across nodule zones, the relative size of nitrogen fixation zones in branched nodules, and the potential for rhizobia to increase their fitness further after the population size reaches a plateau by accumulating polyhydroxybutyrate (PHB), an energy source that supports their fitness in the soil (Ratcliff et al., 2008; Trainer et al., 2010).

### The proportion of branched nodules can be used as a proxy for plant benefit

The use of root nodule count as a metric of symbiotic effectiveness does not account for the size or shape of the nodules. We observed that rhizobia isolates altered plant-level traits, such as nodule number and the proportion of branched nodules. In fact, these two metrics are negatively correlated. Further, we found that nodule branching propensity was strongly correlated with plant benefits, whereas nodule number was not. Our results regarding higher plant benefit with a higher proportion of branched nodules align with Cai et al. (2018), who showed that plants with more branched nodules tend to exhibit an increase in nitrogenase activity per plant and per nodule (Cai et al., 2018). Histological analysis of alfalfa and pea branched nodules has revealed a large nitrogen fixation zone (Lynd and Ansman, 1990; Serova et al., 2019). Thus, our results suggest that the proportion of branched nodules could be used as a proxy for symbiotic investment/benefit that does not require nodule drying, and allows sample processing for measuring microbial communities via sequencing of whole genomes or amplicons (Oono et al., 2011; Brockhurst and Koskella, 2013; Allito et al., 2021).

### Both host and rhizobial genetics influence branching propensity

Our results align with previous studies showing that both the isolate identity and plant host can contribute to individual nodule branching and branching frequency. For instance, Sagan and Gresshoff found that *Rhizobium leguminusarum* strain MSDJ1243 increased the number of individual nodule branches in *Pisum sativum* cultivar. Using the natural genetic diversity of rhizobial isolates, Laguerre et al. found that a subset of *R. leguminosarum* induces larger nodules that exhibit a branched shape and a lower number of nodules (Laguerre et al. 2007). This aligns with the positive correlation between nodule dry biomass and branching reported by Sagan and Gresshoff (Sagan and Gresshoff, 1996). Furthermore, the isolates that induced larger and branched nodules in the latter study have a NodD gene that matches that of isolate MSDJ1243 in the former study. Nod genes have also been associated with nodule branching in *M. truncatula* where rhizobia that overexpress nod genes by carrying a plasmid with extra copies of genes tend to form more branches in *M. truncatula* R108 (Schultze et al., 1992; Cai et al., 2018). The only plant genes known to influence nodule morphology are the MtNFH1and *coch* in *Pisum sativum* (Ferguson and Reid, 2005; Cai et al., 2018). It is unclear whether branches form because the nodules are larger and tend to branch as they age, or if certain strains directly promote branching during early development (D’Amours et al., 2024). We found small, and potentially young, branched nodules, suggesting that branching may occur independently of nodule age in some cases. Further studies are needed to clarify which host and strain genes are associated with the induction of higher nodule branching.

### Nodule branching could play an adaptive role in mutualism evolution

We found that both host and rhizobial genetic variation contribute to nodule branching and that nodule branching is linked to both rhizobial and plant fitness. Despite our observations being limited to host investment, as we worked with single-isolate inoculations, these results suggest that nodule branching could influence mutualism evolution and stability through the reward/sanction mechanism (Wendlandt et al., 2019; Porter et al., 2024). For instance, if nodules containing more beneficial rhizobia receive more rewards from their hosts, they would therefore be more likely to form branches, which could both enable additional areas for nitrogen fixation and increase rhizobial population size. This aligns with previous studies using other nodule traits, such as nodule size or nodule fresh weight (Kiers et al., 2003; Westhoek et al., 2021). One mixed inoculation study found little evidence that individual nodule branching was correlated with plant benefit when inoculating three *M. truncatula* with a mix of *Sinorhizobium* (Heath and Tiffin, 2009), but this study was underpowered to assess this question and used strains with small differences in nitrogen fixation that could be more nuanced than hosts can detect (Oono et al., 2011). While further multi-strain inoculations are needed to assess if branching is relevant for sanctioning independently of size, recent data using pooled sequencing of nodules show that strains inhabiting large, branched nodules are enriched for beneficial strains relative to pools of small, less branched nodules (Burghardt et al., 2025). Overall, nodule branching could provide an additional axis of trait variation, potentially altering the dynamics of legume-rhizobium evolution (Sachs et al., 2018).

### Broader Implications for evaluating nodule shape metrics for managing beneficial microbes

Overall, our results suggest that nodule branching traits should be included in future studies focusing on the influence of rhizobial and host genes on symbiosis. For instance, measuring nodule traits within species using diversity panels (Burghardt et al., 2025) could reveal allelic variation associated with nodule traits, and assessments of how host and rhizobial benefit traits differ across contrasting nodule development strategies or environmental contexts (D’Amours et al., 2024). Alongside other metrics, the proportion of branched nodules could add to efforts to improve nitrogen fixation and thus decrease reliance on synthetic nitrogen fertilizers (Herridge et al., 2008; Gopalakrishnan et al., 2015; Mendoza-Suárez et al., 2021). More broadly, our results suggest that applying evolutionary ecology perspectives to the development of symbiotic organs can provide novel insights and new traits even in well-studied model systems like legumes and rhizobia.

## Abbreviations

ANOVA: analysis of variance
CFU: colony forming units
DI: de-inonized
OD600: optical density
PHB: polyhydroxybutyrate

## Supplementary data

The following supplementary data are available at JXB online.

Table S1. ANOVA results for linear models of individual nodule traits.

Table S2. ANOVA results for linear models of host plant traits.

## Acknowledgements

We thank additional members of the Burghardt lab for assisting in this experiment’s setup, maintenance, and take-down. Huck Genomics Core Facility (RRID: SCR_023645) for library prep and sequencing of rhizobial isolates. Russel E Larson Agricultural Station for hosting the sourced field locations, Tyson building’s greenhouse manager Scott Diloreto for providing materials and guidance during this experiment.

## Author Contributions

LTB conceived the research idea, designed the experiment, contributed to the analysis, and revised the manuscript. SVG and MAGP also contributed to the experimental design and execution, including the measurement of rhizobial population sizes. AAS assisted with plant care and led the analysis of nodule images. ELP managed and executed the greenhouse experiment, analyzed the data, drafted and led the revision of the manuscript. All authors edited the manuscript.

### Generative Artificial Intelligence (AI) statement

No generative AI was used in the writing and preparation of this article. All data analysis, writing, and interpretation were done by the authors.

## Data Availability

GitHub Repository: github.com/EPaillan/nodule_branching_morphology

## REFERENCES

Allito BB, Ewusi-Mensah N, Logah V, Hunegnaw DK (2021) Legume-rhizobium specificity effect on nodulation, biomass production and partitioning of faba bean (Vicia faba L.). Sci Rep 11: 3678

Barker DG, Pfaff T, Moreau D, Groves E, Ruffel S, Lepetit M, Whitehand S, Maillet F, Nair RM, Journet E-P (2006) Growing M. truncatula: choice of substrates and growth conditions.

Batut J, Boistard P (1994) Oxygen control in Rhizobium. Antonie van Leeuwenhoek 66: 129–150

Brockhurst MA, Koskella B (2013) Experimental coevolution of species interactions. Trends in Ecology & Evolution 28: 367–375

Burghardt L, Sydow P, Sutherland J, Epstein B, Tiffin P (2025) Genetic variation in host selectivity and adaptive strain enrichment within legume-rhizobia symbiosis: processes are host-dependent, far from perfect, and correlate with nodule morphology. 2025.06.16.659998

Burghardt LT (2020) Evolving together, evolving apart: measuring the fitness of rhizobial bacteria in and out of symbiosis with leguminous plants. New Phytol 228: 28–34

Cai J, Zhang L-Y, Liu W, Tian Y, Xiong J-S, Wang Y-H, Li R-J, Li H-M, Wen J, Mysore KS, et al (2018) Role of the Nod Factor Hydrolase MtNFH1 in Regulating Nod Factor Levels during Rhizobial Infection and in Mature Nodules of Medicago truncatula. The Plant Cell 30: 397–414

Camerini S, Senatore B, Lonardo E, Imperlini E, Bianco C, Moschetti G, Rotino GL, Campion B, Defez R (2008) Introduction of a novel pathway for IAA biosynthesis to rhizobia alters vetch root nodule development. Arch Microbiol 190: 67–77

D’Amours E, Bertrand A, Cloutier J, Claessens A, Rocher S, Seguin P (2024) Impact of Sinorhizobium meliloti strains and plant population on regrowth and nodule regeneration of alfalfa after a freezing event. Plant Soil 500: 161–179

Denison RF, Kiers ET (2004) Lifestyle alternatives for rhizobia: mutualism, parasitism, and forgoing symbiosis. FEMS Microbiology Letters 237: 187–193

Dupont L, Alloing G, Pierre O, El S, Hopkins J, Hrouart D, Frendo P (2012) The Legume Root Nodule: From Symbiotic Nitrogen Fixation to Senescence. Senescence. doi: 10.5772/34438

Ferguson BJ, Indrasumunar A, Hayashi S, Lin M-H, Lin Y-H, Reid DE, Gresshoff PM (2010) Molecular Analysis of Legume Nodule Development and Autoregulation. Journal of Integrative Plant Biology 52: 61–76

Ferguson BJ, Reid JB (2005) Cochleata: Getting to the Root of Legume Nodules. Plant Cell Physiol 46: 1583–1589

Gola EM (2014) Dichotomous branching: the plant form and integrity upon the apical meristem bifurcation. Frontiers in Plant Science 5:

Gopalakrishnan S, Sathya A, Vijayabharathi R, Varshney RK, Gowda CLL, Krishnamurthy L (2015) Plant growth promoting rhizobia: challenges and opportunities. 3 Biotech 5: 355–377

Heath KD, Tiffin P (2009) Stabilizing mechanisms in a legume-rhizobium mutualism. Evolution 63: 652–662

Herridge DF, Peoples MB, Boddey RM (2008) Global inputs of biological nitrogen fixation in agricultural systems. Plant Soil 311: 1–18

Hirsch AM (1992) Developmental biology of legume nodulation. New Phytologist 122: 211– 237

Kiers ET, Rousseau RA, West SA, Denison RF (2003) Host sanctions and the legume– rhizobium mutualism. Nature 425: 78–81

Laboratory BN (1954) Abnormal and Pathological Plant Growth: Report of Symposium Held August 3 to 5, 1953. Biology Department, Brookhaven National Laboratory

Laguerre G, Depret G, Bourion V, Duc G (2007) Rhizobium leguminosarum bv. Viciae genotypes interact with pea plants in developmental responses of nodules, roots and shoots. New Phytologist 176: 680–690

Lindström K, Mousavi SA (2019) Effectiveness of nitrogen fixation in rhizobia. Microb Biotechnol 13: 1314–1335

Łotocka B, Kopcińska J, Skalniak M (2012) Review article: The meristem in indeterminate root nodules of Faboideae. Symbiosis 58: 63–72

Lu P, Werb Z (2008) Patterning Mechanisms of Branched Organs. Science 322: 1506–1509

Lynd JQ, Ansman TR (1990) Soil constraints with distinctive coralloid nodulation and nitrogen fixation of “Mecca” alfalfa. Journal of Plant Nutrition 13: 77–94

Mendoza-Suárez M, Andersen SU, Poole PS, Sánchez-Cañizares C (2021) Competition, nodule occupancy, and persistence of inoculant strains: key factors in the Rhizobium-legume symbioses. Front Plant Sci 12: 690567

Moran NA (2007) Symbiosis as an adaptive process and source of phenotypic complexity. Proceedings of the National Academy of Sciences 104: 8627–8633

Oldroyd GED (2013) Speak, friend, and enter: signalling systems that promote beneficial symbiotic associations in plants. Nat Rev Microbiol 11: 252–263

Oldroyd GED, Murray JD, Poole PS, Downie JA (2011) The Rules of Engagement in the Legume-Rhizobial Symbiosis. Annu Rev Genet 45: 119–144

Oono R, Anderson CG, Denison RF (2011) Failure to fix nitrogen by non-reproductive symbiotic rhizobia triggers host sanctions that reduce fitness of their reproductive clonemates. Proc Biol Sci 278: 2698–2703

Porter SS, Dupin SE, Denison RF, Kiers ET, Sachs JL (2024) Host-imposed control mechanisms in legume–rhizobia symbiosis. Nat Microbiol 9: 1929–1939

Ratcliff WC, Kadam SV, Denison RF (2008) Poly-3-hydroxybutyrate (PHB) supports survival and reproduction in starving rhizobia. FEMS Microbiology Ecology 65: 391– 399

Roux B, Rodde N, Jardinaud M, Timmers T, Sauviac L, Cottret L, Carrère S, Sallet E, Courcelle E, Moreau S, et al (2014) An integrated analysis of plant and bacterial gene expression in symbiotic root nodules using laser-capture microdissection coupled to RNA sequencing. The Plant Journal 77: 817–837

Sachs JL, Quides KW, Wendlandt CE (2018) Legumes versus rhizobia: a model for ongoing conflict in symbiosis. New Phytologist 219: 1199–1206

Sagan M, Gresshoff PM (1996) Developmental mapping of nodulation events in pea (Pisum sativum L.) using supernodulating plant genotypes and bacterial variability reveals both plant and Rhizobium control of nodulation regulation. Plant Science 117: 167–179

Schneider CA, Rasband WS, Eliceiri KW (2012) NIH Image to ImageJ: 25 years of image analysis. Nature Methods 9: 671–5

Schultze M, Quiclet-Sire B, Kondorosi E, Virelizer H, Glushka JN, Endre G, Géro SD, Kondorosi A (1992) Rhizobium meliloti produces a family of sulfated lipooligosaccharides exhibiting different degrees of plant host specificity. Proc Natl Acad Sci U S A 89: 192–196

Serova TA, Tsyganova AV, Tikhonovich IA, Tsyganov VE (2019) Gibberellins Inhibit Nodule Senescence and Stimulate Nodule Meristem Bifurcation in Pea (Pisum sativum L.). Front Plant Sci. doi: 10.3389/fpls.2019.00285

Sprent JI, Ardley J, James EK (2017) Biogeography of nodulated legumes and their nitrogen-fixing symbionts. New Phytologist 215: 40–56

Thies JE, Singleton PW, Bohlool BB (1991) Influence of the Size of Indigenous Rhizobial Populations on Establishment and Symbiotic Performance of Introduced Rhizobia on Field-Grown Legumes. Appl Environ Microbiol 57: 19–28

Timmers AC, Soupène E, Auriac MC, de Billy F, Vasse J, Boistard P, Truchet G (2000) Saprophytic intracellular rhizobia in alfalfa nodules. Mol Plant Microbe Interact 13: 1204–1213

Tjepkema JD, Yocum CS (1973) Respiration and oxygen transport in soybean nodules. Planta 115: 59–72

Trainer MA, Capstick D, Zachertowska A, Lam KN, Clark SR, Charles TC (2010) Identification and characterization of the intracellular poly-3-hydroxybutyrate depolymerase enzyme PhaZ of Sinorhizobium meliloti. BMC Microbiology 10: 92

Udvardi M, Poole PS (2013) Transport and Metabolism in Legume-Rhizobia Symbioses. Annu Rev Plant Biol 64: 781–805

Vasse J, de Billy F, Camut S, Truchet G (1990) Correlation between ultrastructural differentiation of bacteroids and nitrogen fixation in alfalfa nodules. J Bacteriol 172: 4295–4306

Vincent JM (1970) A manual for the practical study of root nodule bacteria / J.M. Vincent. Blackwell Scientific

Visick KL, Stabb EV, Ruby EG (2021) A lasting symbiosis: how Vibrio fischeri finds a squid partner and persists within its natural host. Nat Rev Microbiol 19: 654–665

Wendlandt CE, Regus JU, Gano-Cohen KA, Hollowell AC, Quides KW, Lyu JY, Adinata ES, Sachs JL (2019) Host investment into symbiosis varies among genotypes of the legume Acmispon strigosus, but host sanctions are uniform. New Phytologist 221: 446–458

Westhoek A, Clark LJ, Culbert M, Dalchau N, Griffiths M, Jorrin B, Karunakaran R, Ledermann R, Tkacz A, Webb I, et al (2021) Conditional sanctioning in a legume– Rhizobium mutualism. Proceedings of the National Academy of Sciences 118: e2025760118

