## Supplementary Tables S1-S2 for "Nodule branching, size, and symbiosis outcomes shaped by natural genetic variation in rhizobia and alfalfa"

**Supplementary Information**

**Nodule branching, size, and symbiosis outcomes are shaped by natural genetic variation in rhizobial bacteria and alfalfa hosts**

Elizabeth L. Paillan^1, 2^, Alejandra Gil-Polo^1^, Sohini Guha^1^, Andy Swartley^1^, and Liana T. Burghardt^1^

**Table S1:** ANOVA results for linear models of individual nodule traits for Figure 2a-c, respectively.

| **Trait** | **Predictor** | **Df^1^** | **Prop. Variance^2^** | | | **F value^3^** | | ***P* value^4^** | **Model R^2^ adj^5^** |
| --- | --- | --- | --- | --- | --- | --- | --- | --- | --- |
| Log_10_(  Nodule surface area) | **Number of branches** | 3 | | 0.637 | 157.71 | | *4.14×10^-48^**** | | 0.6678 |
|  | Host variety | 2 | | 0.001 | 0.50 | | *0.607* | |  |
|  | **Rhizobial isolate** | 7 | | 0.037 | 3.89 | | *0.001*** | |  |
|  | **Replicate** | 3 | | 0.014 | 3.54 | | *0.02** | |  |
|  | Number of branches*Host variety | 6 | | 0.009 | 1.09 | | 0.37 | |  |
|  | Number of branches*  Rhizobial isolate | 20 | | 0.027 | 1.01 | | 0.46 | |  |
|  | Host*isolate | 14 | | 0.023 | 1.22 | | 0.27 | |  |
|  | Number of branches* Host variety *Rhizobial isolate | 28 | | 0.032 | 0.85 | | 0.69 | |  |
|  | Residual | 164 | | 0.221 | NA | | NA | |  |
| Log_10_(  Rhizobia bacterial colonies) | **Number of branches** | 3 | | 0.183 | 19.07 | | *4.66×10^-11^**** | | 0.2116 |
|  | **Host variety** | 2 | | 0.041 | 6.40 | | *0.002*** | |  |
|  | Rhizobial isolate | 7 | | 0.026 | 1.14 | | 0.34 | |  |
|  | Replicate | 3 | | 0.013 | 1.39 | | 0.25 | |  |
|  | Number of branches *  Host variety | 6 | | 0.016 | 0.86 | | 0.53 | |  |
|  | Residuals | 226 | | 0.721 | NA | | NA | |  |
| Log_10_(  Rhizobia bacterial colonies) | **Nodule surface area** | 1 | | 0.248 | 112.09 | | *1.50×10^-21^**** | | 0.453 |
|  | Number of branches | 3 | | 0.013 | 1.95 | | 0.12 | |  |
|  | **Host variety** | 2 | | 0.032 | 7.13 | | *0.001** | |  |
|  | Rhizobial Isolate | 7 | | 0.247 | 1.59 | | 0.14 | |  |
|  | Replicate | 3 | | 0.006 | 0.95 | | 0.41 | |  |
|  | **Nodule surface area***  **Number of branches** | 3 | | 0.171 | 25.73 | | *2.3×10-^14^**** | |  |
|  | Residuals | 228 | | 0.505 | NA*^6^ | | NA | |  |

^1^Df = degrees of freedom.

^2^Prop. variance = Proportion of total variance explained by each predictor

^3^F value = F statistic from the ANOVA model.

^4^Pval = significance level of the predictor effect (*=Pval<0.05;**= Pval< 0.01; ***= Pval< 0.001)

^5^Model R^2^ adjusted

^6^NA = not applicable.

**Table S2:** ANOVA results for linear models of host plant traits in Figure 3 a-c, respectively.

| **Trait** | **Predictor** | **Df^1^** | **Prop.**  **Variance^2^** | | **F value^3^** | | ***P* value^4^** | **Model R^2^ adj^5^** |
| --- | --- | --- | --- | --- | --- | --- | --- | --- |
| Mean host biomass (g) | **Mean proportion branched nodules** | 1 | | 0.154 | 118.82 | *1.13×10^-22^**** | | 0.530 |
|  | **Host variety** | 2 | | 0.144 | 55.51 | *1.7×10^-20^**** | |  |
|  | **Rhizobial isolate** | 121 | | 0.391 | 2.49 | *1.27×10^-9^**** | |  |
|  | Proportion branched*Host variety | 2 | | 0.005 | 1.75 | 0.1754 | |  |
|  | residual | 236 | | 0.306 | NA | NA | |  |
| Mean nodule count | **Mean proportion branched nodules** | 1 | | 0.121 | 80.92 | *7.89×10^-17^**** | | 0.458 |
|  | **Host variety** | 2 | | 0.080 | 26.85 | *3.11×10^-11^**** | |  |
|  | **Rhizobial isolate** | 121 | | 0.442 | 2.45 | *2.28×10^-9^**** | |  |
|  | Proportion branches* Host variety | 2 | | 0.002 | 0.67 | *0.5123* | |  |
|  | residual | 236 | | 0.353 | NA^6^ | NA | |  |
| Proportion of branched nodules | **Host variety** | 2 | | 0.006 | 4.12 | *0.02** | | 0.454 |
|  | **Rhizobial Isolate** | 121 | | 0.261 | 3.03 | *3.19 × 10^-20^**** | | |
|  | Replicate | 7 | | 0.006 | 1.18 | 0.31 | |  |
|  | Host variety*Rhizobial isolate | 239 | | 0.182 | 1.07 | 0.25 | |  |
|  | Residual | 769 | | 0.546 | NA | NA | |  |

^1^Df = degrees of freedom.

^2^Prop. variance = Proportion of total variance explained by each predictor

^3^F value = F statistic from the ANOVA model.

^4^Pval = significance level of the predictor effect (*=Pval<0.05;**= Pval< 0.01; ***= Pval< 0.001)

^5^Model R^2^ adjusted

^6^NA = not applicable.
